# Expression, Purification, and Crystallization of Recombinant Human ABL-1 Kinase for Structure-Based Drug Screening Applications

**DOI:** 10.1101/2025.06.02.657364

**Authors:** Ayca Irgit, Halilibrahim Ciftci, Hasan Demirci

## Abstract

Abelson-1 (ABL-1) is a non-receptor tyrosine kinase that plays essential roles in various cellular processes, including proliferation, survival, differentiation and its kinase activity is tightly regulated. The dysregulated ABL-1 kinase activity is linked to disease pathogenesis like Chronic Myeloid Leukemia (CML), where the BCR::ABL-1 fusion oncoprotein drives oncogenic signaling. Due to its central role in CML pathogenesis, understanding the structure of ABL-1 is crucial for the effective management of the disease and drug development studies. This study focuses on optimizing the expression, purification and crystallization of the recombinant human ABL-1 kinase domain for its structural analysis via X-ray crystallography and structure-based drug screening applications. The human ABL-1 kinase domain, fused with a SUMO-tag, was expressed in Escherichia coli Rosetta2™ BL21 using the pET28(a)+ expression vector. The ABL-1 aggregates seen under native culture conditions were successfully solubilized by mild ionic detergent sarkosyl. After obtaining soluble expression of the protein, Ni-NTA affinity chromatography was performed and high tiled of purified ABL-1 was obtained. The 6X-His-SUMO-tag of purified ABL1 was cleaved by ULP1 protease. The recombinant ABL-1 was subsequently used in crystallization trials for enlightening structural features of ABL-1 that could guide the development of novel therapeutics and drug screening platforms targeting ABL-1 in CML.

## 1. Introduction

Abelson-1 (ABL-1) is a non-receptor tyrosine kinase protein belonging to the Abelson (ABL) family and plays a crucial role in cellular processes such as proliferation, survival, differentiation, stress responses, and cytoskeletal reorganization [1-3]. ABL-1 has a complex structure that is key to its diverse cellular functions. It consists of an N-terminal cap region, Src Homology-2 (SH2) and Src Homology-3 (SH3) domains, a bilobed kinase domain (KD), also referred to as the Src Homology-1 (SH1) domain, a Proline-rich motif (PxxP), and a long C-terminal tail, a DNA-binding domain, a G-actin binding domain, and both Nuclear Localization and Nuclear Export Signals [4,5]. The kinase domain of ABL-1 exhibits kinase activity and promotes the protein’s autophosphorylation upon ATP binding [6]. Under normal cellular conditions, ABL-1 remains inactive [7] and its kinase activity is strictly regulated to prevent abnormal kinase activity [8]. Kinase activation usually occurs through autophosphorylation or protein interactions in response to specific cellular signals [7-9]. Abnormal ABL-1 kinase activity has been linked to the cellular abnormalities and disease pathogenesis, including Chronic Myeloid Leukemia (CML) [8].

CML is a hematologic malignancy classified as a myeloproliferative neoplasm, characterized by the unregulated proliferation of myeloid granulocytic cells in the bone marrow [10-12]. The development of CML is mainly driven by the BCR::ABL-1 oncoprotein, which arises from the reciprocal translocation of *ABL1* gene on chromosome 9 with the *Breakpoint Cluster Region (BCR)* gene on chromosome 22 t(9;22)(q34;q11) in bone marrow cells [13-15]. The BCR::ABL-1 oncoprotein has constitutive tyrosine kinase activity due to the loss of regulatory domains caused by the translocation [13, 14]. The constitutive activity of the BCR::ABL-1 oncoprotein triggers oncogenic signaling pathways that lead to uncontrolled cell proliferation and resistance to apoptosis [16, 17]. Tyrosine kinase inhibitors (TKIs) that target ABL-1 kinase are crucial in the treatment of CML [18] by inhibiting the constitutive kinase activity of the BCR::ABL-1 fusion protein [19, 20]. The structural studies that focus on the enlightening structural features of the ABL-1 kinase are crucial for the development of novel TKIs and therapeutic approaches for the effective management of CML [20].

This study aims the optimization of the expression, purification and crystallization of recombinant human ABL-1 in *Escherichia coli* (*E. coli*) for drug screening applications. This study presents an efficient method for the recombinant ABL-1 production, enabling its use in structural studies, drug development, and the investigation of biochemical mechanisms underlying kinase-targeted therapeutics.

## 2. Materials and Methods

### 2.1. Plasmid Construct

The kinase domain of human ABL-1, spanning residues 229–503 (PDB: 2HYY), was selected and modified with an N-terminal SUMO tag to enhance protein solubility. The pET-28a(+) bacterial expression vector, which includes a 6X-His tag and kanamycin resistance as the selection marker, was chosen as the expression vector **(Figure 1)**. The corresponding sequence of SUMO-tagged ABL-1 kinase was synthesized and cloned into the pET28a(+) expression vector by GenScript, USA.

**Figure 1:**
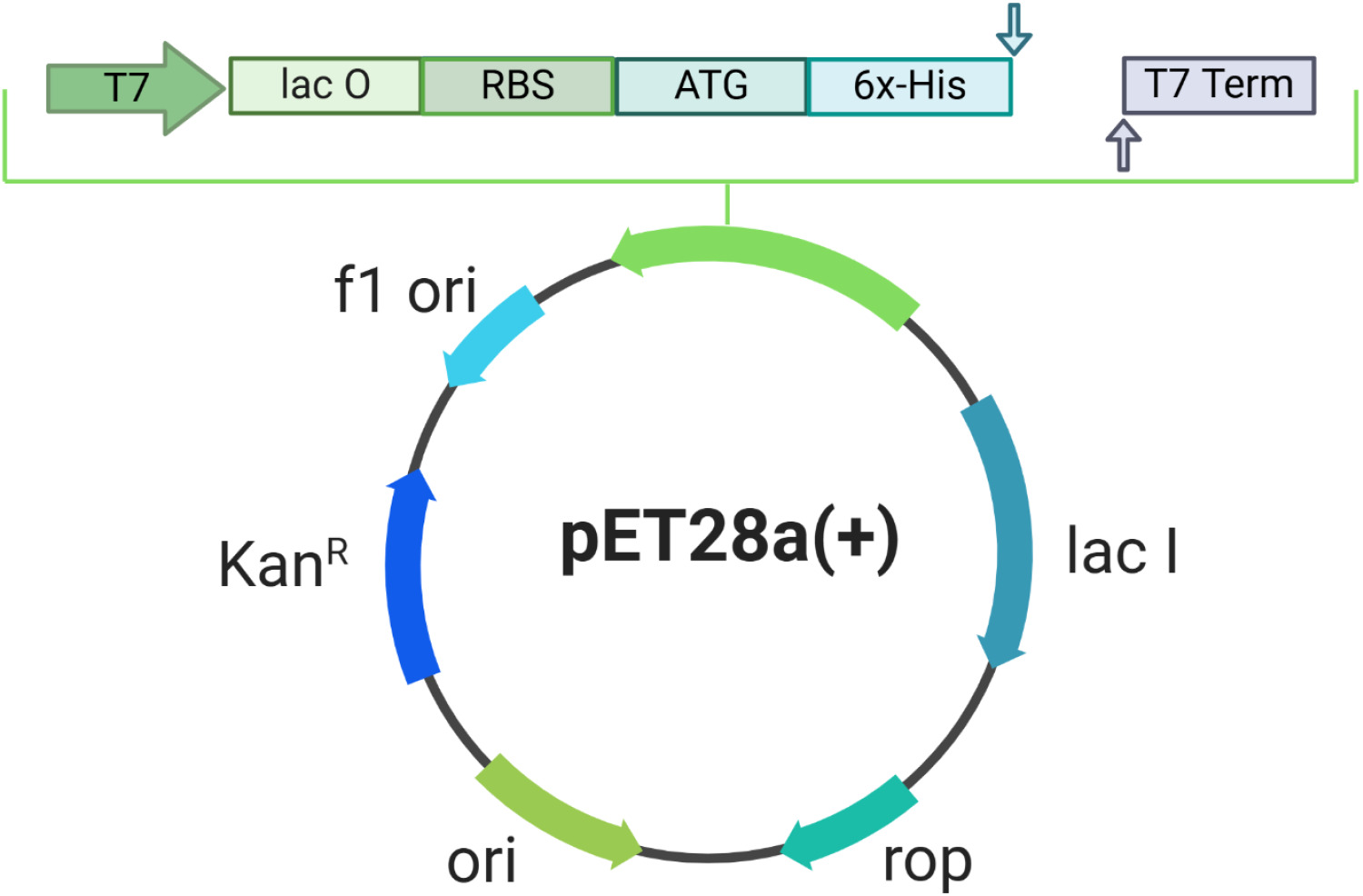
Schematic overview showing the construction of the bacterial expression vector pET-28a(+). Created via BioRender.

### 2.2. Recombinant ABL-1 Expression

Plasmids were transformed into *E. coli* Rosetta2™ BL21 competent cells using the heat shock method by applying heat at 42°C for 45 seconds. The transformed cells were grown on Luria-Bertani (LB) agar plates containing 50 μg/mL kanamycin (Cat#KB0286, Bio Basic, Canada) and 35 μg/mL chloramphenicol (Cat#CB0118, Bio Basic, Canada) at 37°C overnight. A single colony from agar plate was selected and inoculated into 10 mL of LB medium containing 50 μg/mL kanamycin and 35 μg/mL chloramphenicol, then incubated at 37°C and 110 rpm overnight.

Then, 0.4 mM Isopropyl-ß-D-thiogalactopyranoside (IPTG) (Cat#I2481C, GoldBio, USA) was added to the culture and incubated at 18°C and 110 rpm for 17 hours. After incubation, cells were harvested by centrifugation at 6,000 rpm for 5 minutes. The resulting cell pellets were resuspended in 750 μL lysis buffer (500 mM NaCl, 50 mM Tris-HCl, 20 mM Imidazole, %5 glycerol, %0.1 Triton 100-X, pH: 7.5). Then, the cell culture was sonicated until the viscosity was reduced. Following sonication, the suspension was centrifuged at 10,000 rpm for 10 minutes. After centrifugation, a 40 μL sample was taken from the supernatant for SDS-PAGE analysis. The remaining supernatant was transferred to a new, clean tube, and 50 μL of Ni-NTA beads were added, and incubated at 4°C, overnight. After incubation, protein-bead mixture was washed with His A Buffer (200 mM NaCl, 20 mM Tris-HCl, 20 mM Imidazole, pH: 7.5) and the protein was eluted with His B Buffer (200 mM NaCl, 20 mM Tris-HCl, 250 mM Imidazole, pH: 7.5). All samples were analyzed by SDS-PAGE.

### 2.3 Soluble ABL-1 Expression

The soluble expression of ABL-1 was checked under the detergent treatment. Two different detergents urea and sarkosyl were used to test their effects on the solubility of ABL-1 aggregates. 10 mL of two cell cultures were prepared according to the described method and the pellets were lysed with the lysis buffers listed in the **Table 1**, followed by sonication. Following sonication, the described method was applied for both samples, and all fractions were analyzed by SDS-PAGE.

**Table 1:** The lysis buffers and compositions that were used in the solubility test of ABL-1.

| Lysis Buffer # | Composition |
| --- | --- |
| Lysis Buffer#1 | 500 mM NaCl, 50 mM Tris-HCl, 5% glycerol, 0.1% Triton X-100, 8 M Urea (pH 8.0) |
| Lysis Buffer#2 | 500 mM NaCl, 50 mM Tris-HCl, 20 mM imidazole, 5% glycerol, 0.1% Triton X-100, 0.5% Sarkosyl (pH 7.5) |

### 2.4. ABL-1 Purification

For the production of 6X His- and SUMO-tagged ABL-1 kinase protein, 50 mL of pre-culture LB medium containing 50 μg/mL kanamycin and 35 μg/mL chloramphenicol was prepared and then incubated overnight at 37°C and 110 rpm. After incubation, the 50 mL pre-culture was scaled up to 1 L of LB medium containing 50 μg/mL kanamycin and 35 μg/mL chloramphenicol and grown at 37°C, 110 rpm. When OD600 reaches ∼0.8, 0.4 mM IPTG was added to the culture and incubated at 18°C and 110 rpm for 17 hours.

Following incubation, cells were harvested by centrifugation at 3,500 rpm for 45 minutes. The resulting cell pellets were resuspended in 50 mL lysis buffer (500 mM NaCl, 50 mM Tris-HCl, 20 mM Imidazole, %5 glycerol, %0.1 Triton 100-X, %0.5 sarkosyl; pH: 7.5). Then, the cell culture was sonicated at 60% power for 20 seconds, repeated five times, until the viscosity was reduced. Following sonication, the suspension was ultracentrifuged at 35,000 rpm and 4°C for 1 hour by using the Ti-45 rotor (Beckman, USA). The supernatant was then transferred into a 3 kDa dialysis membrane and dialyzed overnight at 4°C against 2 L of dialysis buffer (200 mM NaCl, 20 mM Tris-HCl, pH 7.5). After dialysis, the protein sample was filtered through a 0.22-µm hydrophilic polyethersulfone (PES) membrane filter (Cat# SLGP033NS, Merck Millipore, USA).

For the purification of 6XHis-SUMO-tagged ABL-1 kinase protein, Ni-NTA column (Qiagen, Venlo, Netherlands) was used. First, the column was washed with 1 M Imidazole, followed by equilibration with His A buffer (200 mM NaCl, 20 mM Tris-HCl, 20 mM Imidazole, pH: 7.5). After equilibration, the protein sample was loaded onto the column and untagged proteins were collected in flowthrough. Next, the column was washed with the wash buffer (200 mM NaCl, 20 mM Tris-HCl, pH: 7.5) to remove nonspecifically bound proteins. The target protein was eluted using His B buffer (200 mM NaCl, 20 mM Tris-HCl, 250 mM imidazole, pH 7.5) and collected as the elution sample. All collected fractions were analyzed by SDS-PAGE. The elution sample was concentrated 20-fold using a 10 kDa Amicon filter (Merck Millipore, Germany). The concentrated protein sample was then aliquoted, flash-frozen in liquid nitrogen, except one of the aliquot that was used in 6X-His-SUMO tag cleavage trial, and stored at -80°C for crystallization studies.

### 2.5. 6X-His-SUMO-Tag Removal by ULP1 Enzymatic Cleavage

For the cleavage of the 6X-His-SUMO tag from purified ABL-1, a cleavage trial with the ULP1 enzyme was performed at three different incubation temperatures for different durations. The ULP1 enzyme was added to the protein sample at a 1:100 ratio and incubated at 4°C, 25°C, and 37°C. Protein-enzyme mixture was incubated overnight, with samples taken after 1 hour, 2 hours, 4 hours and after overnight incubation for each temperature. The samples were analyzed by SDS-PAGE to check 6X-His-SUMO tag cleavage.

### 2.6. Crystallization and Structural Studies

The crystallization of ABL-1 protein was carried out using the microbatch under oil method in 72-well Terasaki crystallization plates (Cat#654180, Greiner Bio-One, Austria) as previously described [21]. In this process, 0.83 µL of the ABL-1 protein sample was combined with 0.83 µL of approximately 3000 commercially available sparse matrix and grid screen crystallization cocktail solutions **(Table 2)** [21]. 16.6 µL of paraffin oil (Cat#ZS.100510.5000, ZAG Kimya, Turkey) was added to seal each well and the plates were stored at 4°C. The Terasaki plates were regularly controlled under the light microscope to monitor crystal formation. The observed crystals were analyzed at ambient temperature using a Rigaku’s XtaLAB Synergy Flow XRD that is equipped with CrysAlisPro 1.171.42.35a software.

**Table 2:** Crystallization conditions used for crystal screening.

| Hampton Research |  |  |
| --- | --- | --- |
| Natrix (HR2-116) #1-48 | Natrix 2 (HR2-117) #1-48 | Index (HR2-144) #1-96 |
| MembFac (HR2-114) #1-48 | PEG/Ion Screen (HR2-126) #1-48 | PEG/Ion 2 Screen (HR2-098) #1-48 |
| SaltRx 1 (HR2-107) #1-48 | SaltRx 2 (HR2-109) #1-48 | Quik Screen (HR2-221) #1-24 |
| Ionic Liquid Screen (HR2-214) #1-24 | Crystal Screen Cryo (HR2-122) #1-50 | Crystal Screen 2 Cryo (HR2-121) #1-48 |
| Crystal Screen (HR2-110) #1-50 | Crystal Screen 2 (HR2-112) #1-48 | Crystal Screen Lite (HR2-128) |
|  |  | #1-50 |
| PEGRx 1 (HR2-082) #1-48 | PEGRx 2 (HR2-084) #1-48 | Grid Screen PEG 6000 (HR2-213) #1-24 |
| Grid Screen Sodium Chloride (HR2-219) #1-24 | Grid Screen Sodium Malonate (HR2-247) #1-24 | Grid Screen Ammonium Sulfate (HR2- 211) #1-24 |
| Grid Screen MPD (HR2-215) #1-24 | Grid Screen PEG/LiCl (HR2-217) #1-24 |  |
| **Molecular Dimensions** |  |  |
| Wizard Classic 1 (MD15-W1-T) #1-48 | Wizard Classic 2 (MD15-W2-T) #1-48 | Wizard Classic 3 (MD15-W3-T) #1-48 |
| Wizard Classic 4 (MD15-W4-T) #1-48 | JCSG-plus™ (MD1-37) #1-96 | HELIX™ (MD1-68) #1-96 |
| MIDASplus™ (MD1-106) #1-96 | NR-LBD™ (MD1-24) #1-48 | NR-LBD™ Extension (MD1-26) #1-48 |
| ProPlex™ (MD1-38) #1-96 | The PGA Screen™ (MD1-50) #1-96 | Morpheus® (MD1-46) #1-96 |
| Structure Screen 1 (MD1-01) #1-50 | Structure Screen 2 (MD1-02) #1-50 | PACT premier™ (MD1-29) #1-96 |
| Stura FootPrint Screen (MD1-20) #1-48 | MultiXtal (MD1-65) #1-48 | MacroSol™ (MD1-22) #1-48 |
| 3D Structure Screen (MD1-13) #1-48 | Wizard Cryo 1 (MD15-C1-T) #1-48 | Wizard Cryo 2 (MD15-C2-T) #1-48 |
| Clear Strategy™ Screen I (MD1-14) #1-245 at different pHs 4.5, 5.5, 6.5, 7.5, 8.5 | Clear Strategy™ Screen II (MD1-15) #1-245 at different pHs 4.5, 5.5, 6.5, 7.5, 8.5 | Wizard Precipitant Synergy Screen (MD15-PS-T) #1-192 |
| **Jena Bioscience** |  |  |
| JBScreen Nuc-Pro 1 (CS-181) #1-24 | JBScreen Nuc-Pro 2 (CS-181) #1-24 | JBScreen Nuc-Pro 3 (CS-181) #1-24 |
| JBScreen Nuc-Pro 4 (CS-181) #1-24 |  |  |
| **NeXtal Biotechnologies** |  |  |
| NeXtal Protein Complex Suite (130715) #1-96 |  |  |

## 3. Results

### 3.1. The Recombinant Expression of Soluble ABL-1

The kinase domain of human ABL-1 protein was successfully expressed recombinantly in *E. coli* Rosetta2™ BL21. In the SDS-PAGE analysis, protein bands at approximately 42 kDa corresponding to the expected molecular weight of human ABL-1 kinase domain (29 kDa) fused with the 13 kDa SUMO-tag were observed **(Figure 2)**. Recombinant expression of human ABL-1 kinase was achieved under the tested culture conditions, using 0.04 mM IPTG induction for 17 hours at 18 °C. Despite successful expression of ABL-1, the majority of the protein expression was observed as insoluble fraction, accumulating in the pellet rather than the soluble protein fraction in the supernatant **(Figure 2)**.

**Figure 2:**
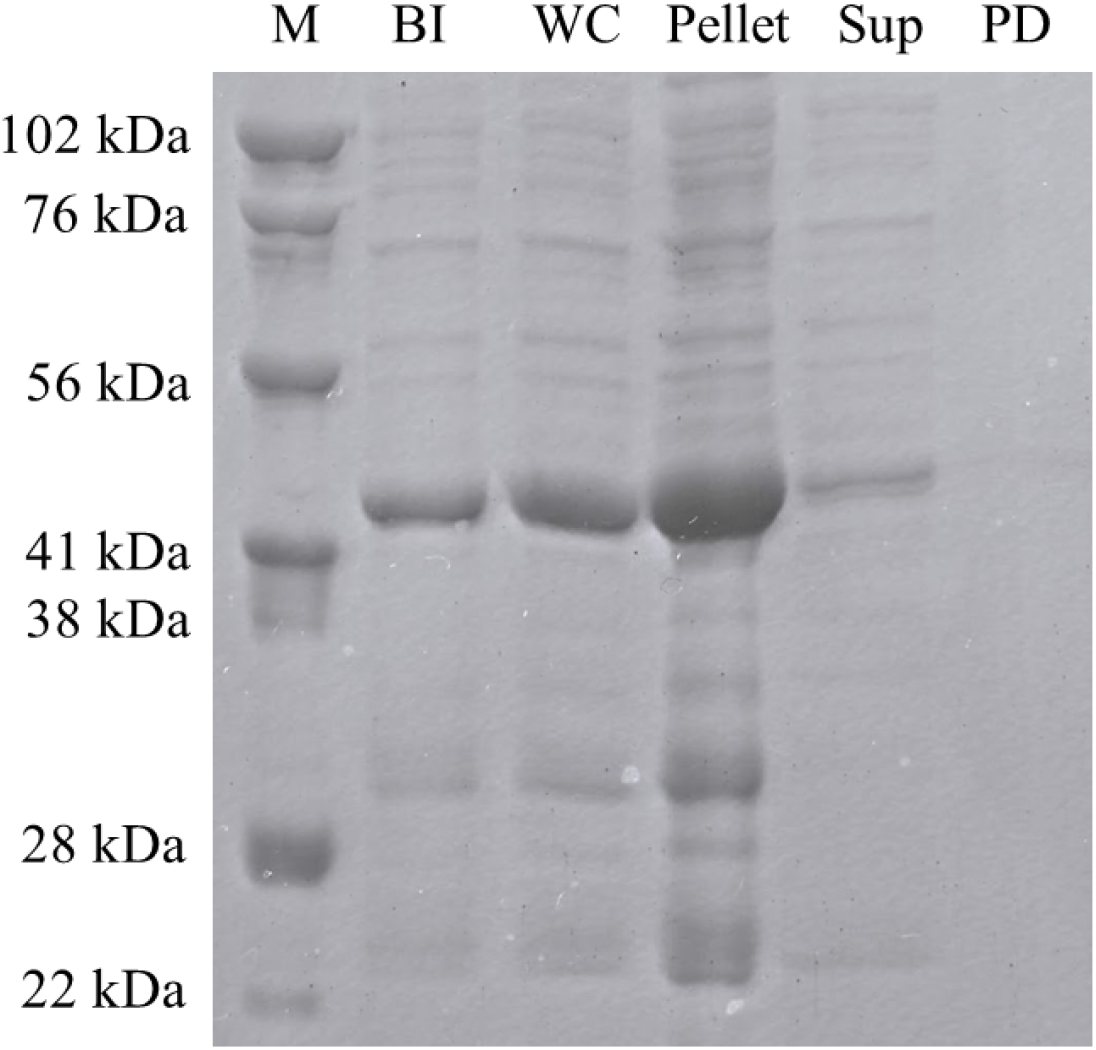
The SDS-PAGE analysis of the recombinant ABL-1 expression in *E. coli* Rosetta-II. M: Marker, BI: Before Induction, WC: Whole Cell Lysate, Sup: Supernatant, PD: Pull down sample.

To achieve soluble ABL-1 expression, lysis buffers with different detergent compositions including urea and sarkosyl were tested. Under both of the urea and sarkosyl treatments, ABL-1 was observed in the supernatant as soluble protein fractions and in pull-down as interacting with Ni-NTA beads **(Figure 3)**. Due to its non-denaturing properties, sarkosyl-containing lysis buffer was used in the subsequent protein production steps.

**Figure 3:**
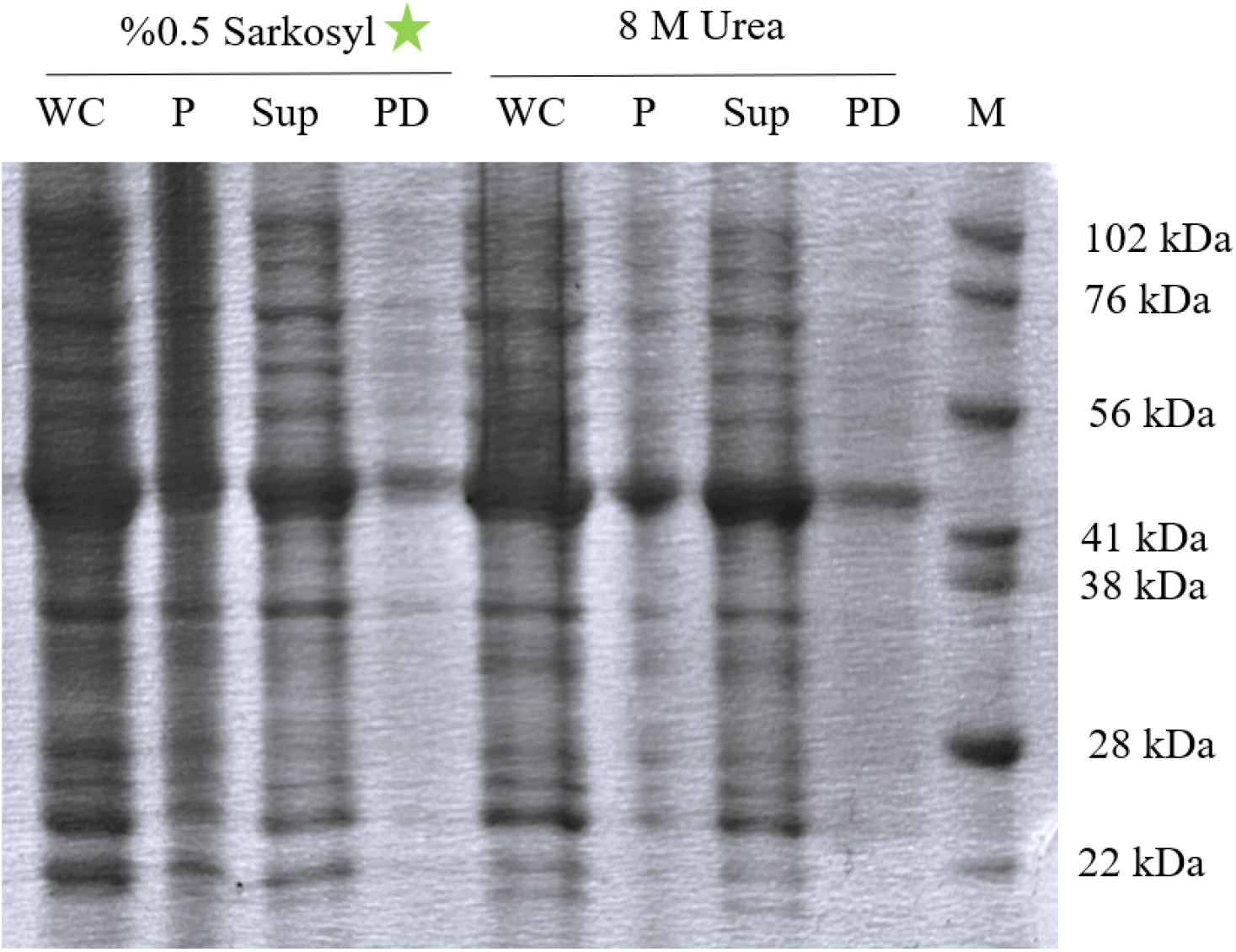
The SDS-PAGE analysis of soluble ABL-1 expression using different lysis buffers. M: Marker, WC: Whole Cell Lysate, P: Pellet, Sup: Supernatant, PD: Pull down sample.

### 3.3. The Purification of ABL-1

Recombinant ABL-1 was successfully purified using Ni-NTA affinity chromatography. After binding to the resin, the target protein was efficiently eluted with a 250 mM imidazole-containing His-B elution buffer. In the SDS-PAGE analysis of the collected elution fractions a distinct, homogeneous band around 42 kDa, which corresponds to the expected molecular weight of ABL-1 (29 kDa) fused to the 13 kDa SUMO-tag were observed **(Figure 4)**. The elution fractions E3-E7, which contains the highest concentrations of purified protein were then concentrated 20-fold **(Figure 5)** to facilitate subsequent crystallization studies of ABL-1.

**Figure 4:**
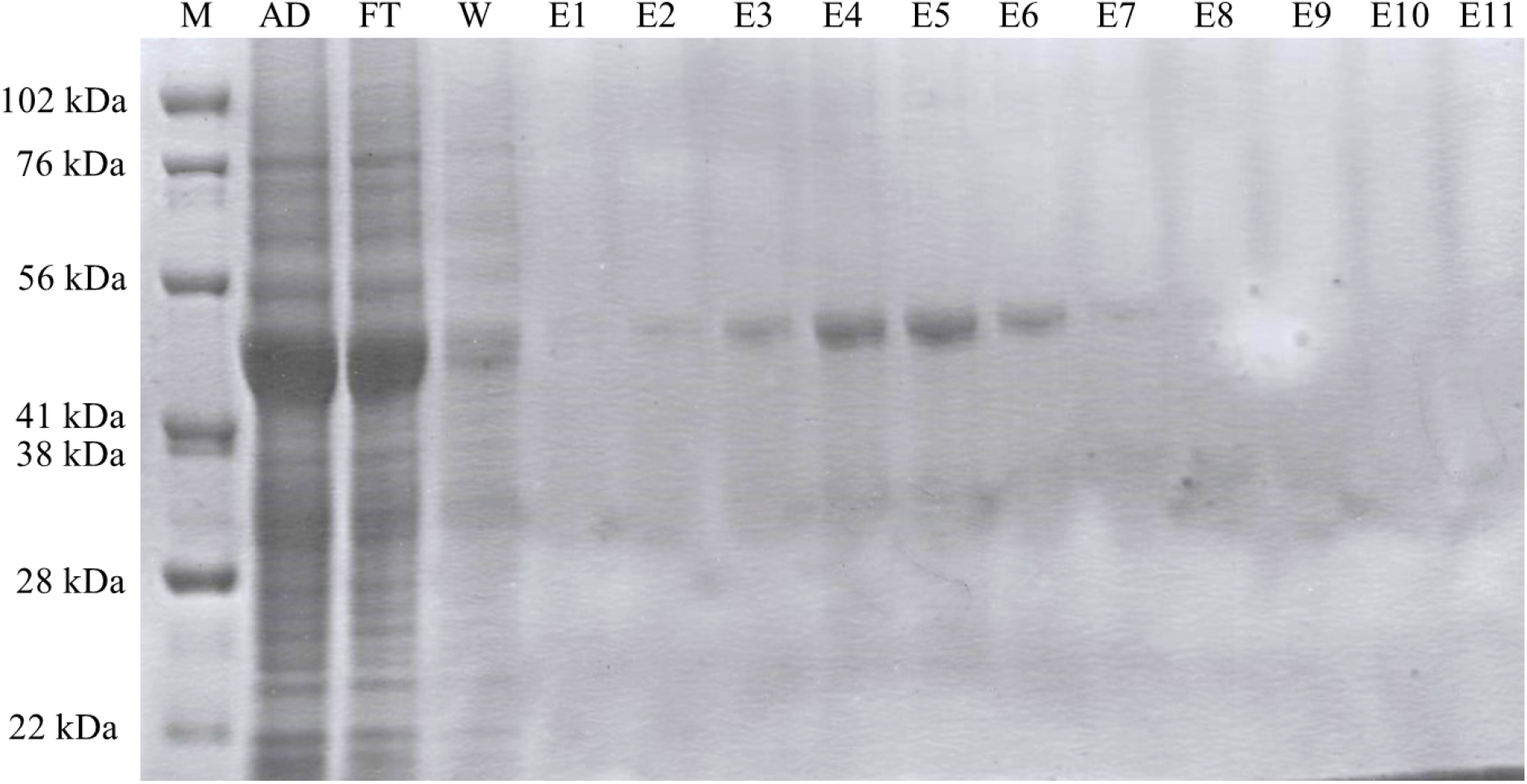
The SDS-PAGE analysis of ABL-1 purification using an NI-NTA column. M: Marker, Sup/AD: Supernatant after dialysis, FT: Flow-through, W: Wash, E1-E11: Elution fragments from 1 to 11.

**Figure 5:**
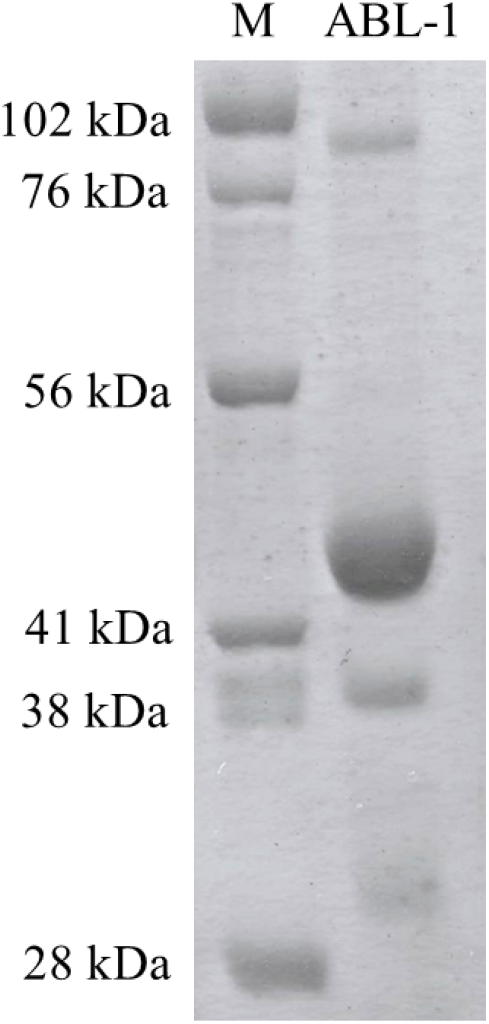
The concentrated ABL-1. M: Marker, ABL-1: Concentrated ABL-1

### 3.4. The Cleavage of ABL-1’s 6X-His-SUMO-tag by ULP1 Enzyme

To obtain tag-free ABL-1, the 6X-His-SUMO-tag was cleaved using the ULP1 enzyme under different incubation conditions. In the SDS-PAGE analysis both the uncut ABL-1 protein (42 kDa, green box) and the cleaved ABL-1 (29 kDa, orange box) with the cleaved SUMO-tag around 13 kDa were observed across all tested conditions **(Figure 6)**. Among all tested conditions, the highest cleavage efficiency was observed at 4°C for higher incubation duration.

**Figure 6:**
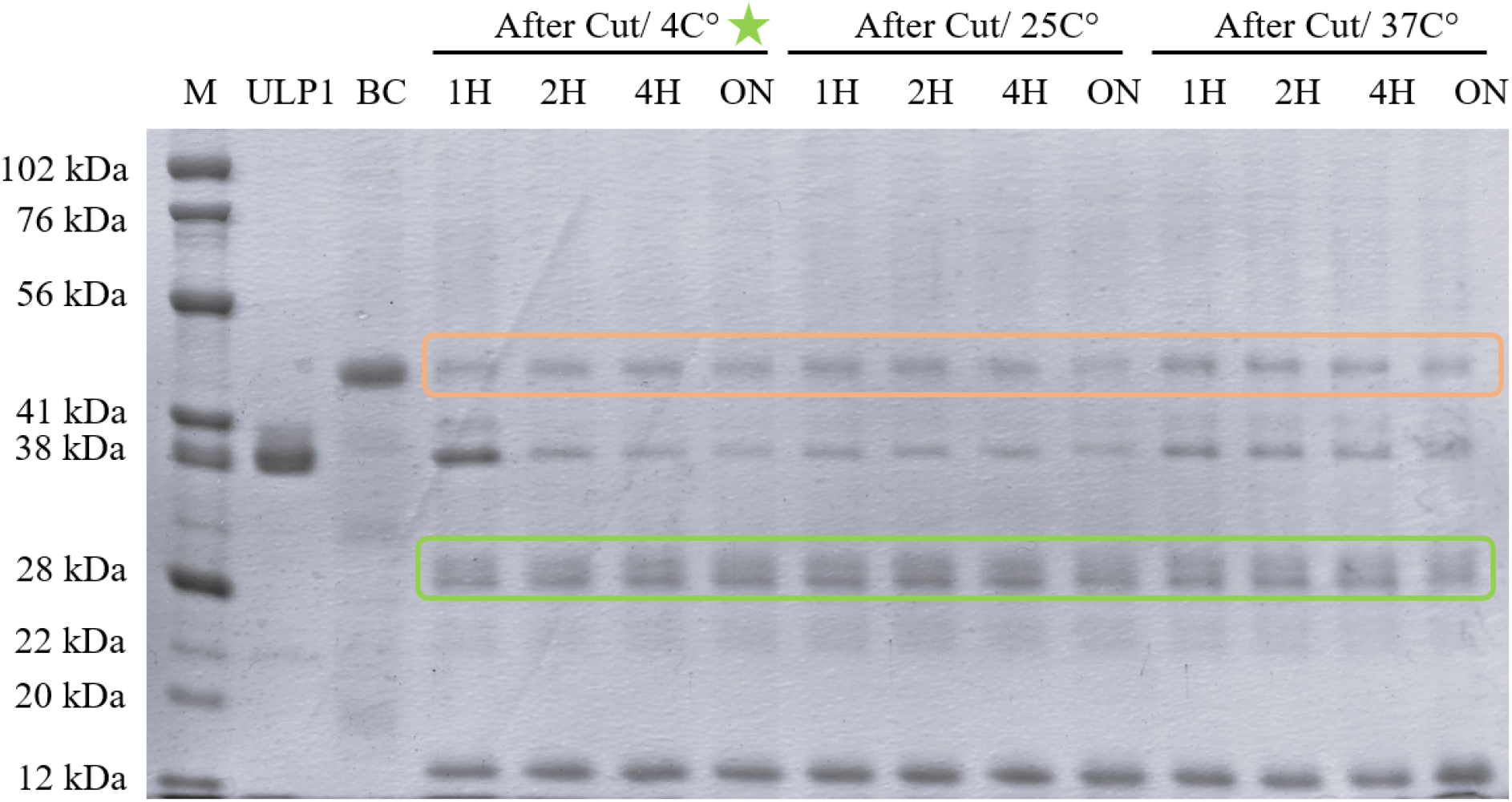
The SDS-PAGE analysis of ABL-1’s 6X-His-SUMO-tag cleavage by ULP1 enzyme at different incubation temperatures and durations. Bands in the green box correspond to the 42 kDa uncut ABL-1 with SUMO-tag, bands in the orange box correspond to the 29 kDa cut ABL-1 without SUMO-tag. M: Marker, BC: Before Cut sample, 1H: The sample taken from after 1 hour incubation, 2H: The sample taken from after 2 hours incubation, 4H: The sample taken from after 4 hours incubation, ON: The sample taken from after overnight incubation.

### 3.5. Crystallization and Structural Analysis of ABL-1

Following the crystallization trials, crystals were observed in several conditions **(Figure 7)**. The monitoring of crystallization plates for crystal formation and their analysis using XRD are ongoing.

**Figure 7:**
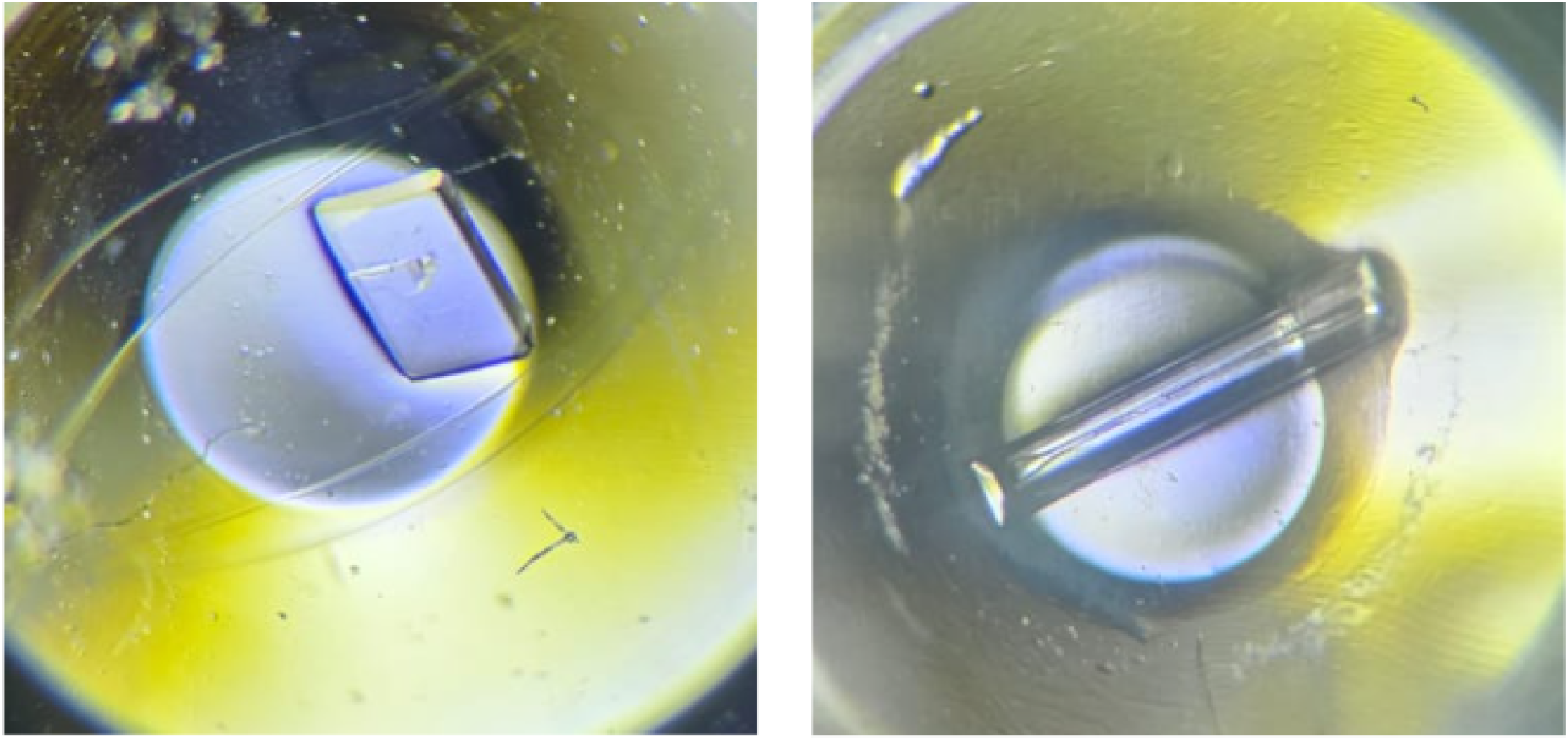
The observed crystals under the light microscope. The crystal was observed under Salt RX I, condition 47 (left). The crystal was observed under Wizard IV condition 44 (right).

## 4. Discussion

Recombinant protein production is the central for the further protein analysis, including structural studies. Therefore, optimizing protein expression and purification strategies is the baseline for most of the protein-related research areas. In this study, we optimized a protein production method for the recombinant human ABL-1 kinase domain to enable its structural analysis via X-ray crystallography and further drug-screening applications.

The human ABL-1 kinase domain was recombinantly expressed using the pET28a(+) expression vector within the *E. coli* Rosetta2™ BL21 host expression system. This vector enhances the simple and efficient purification of the expressed protein through Ni-NTA affinity chromatography by incorporating a 6X-His affinity tag at its N-terminus. To improve the solubility of the expressed ABL-1 protein, a SUMO-tag was integrated at the N-terminus, following the 6X-His tag [22]. Despite the integration of the SUMO-tag, in the SDS-PAGE analysis the majority of the ABL-1 was observed as insoluble fractions in the pellet. This observation suggested that the protein is predominantly expressed as inclusion bodies under native conditions and needs further optimization strategies for ABL-1 production to enhance its solubility and yield.

The soluble expression of ABL-1 was tested using lysis buffers containing varying detergents, urea, and sarkosyl. In both conditions, ABL-1 was detected in the supernatant as soluble fractions. Due to its denaturing properties, the solubilization of the inclusion bodies with urea requires an additional refolding step for proper folding and subsequent purification of the proteins [23]. In contrast, the non-denaturing agent sarkosyl does not require an additional refolding step, allowing the direct purification of the protein [24]. Due to its advantages over urea, sarkosyl was chosen as the solubilizing agent for ABL-1 inclusion bodies. By encapsulating proteins, sarkosyl effectively solubilizes the inclusion bodies [25]. Even at low concentrations (<1%), sarkosyl was efficiently solubilized ABL-1 inclusion bodies. Since it enhances the viscosity, the protein purification becomes more challenging under sarkosyl treatment [25]. So, serial dilutions or dialysis is required to decrease sarkosyl concentration and improve the purification efficiency [25]. Although a low concentration of sarkosyl was used, dialysis was still applied to further enhance purification efficiency.

Following the dialysis, the purification of ABL-1 was successfully performed with minimal impurities or protein degradation via Ni-NTA affinity chromatography and high yield of purified ABL-1 kinase was obtained. After purification, the 6X-His-SUMO tag of ABL-1 was subjected to cleavage by the ULP1 enzyme and the cleaved ABL1 was observed in the gel around 28 kDa.

The produced ABL-1 was used in crystallization trials for further structural analysis and drug-screening applications. Structural studies that aim at enlightening the structural features of ABL-1 are crucial for the management of CML and potential therapeutic advancements for the disease. Structural biology plays a significant role in the development of drug screening platforms by providing atomic-level insights into the three-dimensional architecture of target proteins. Detailed structural information of ABL-1 enables the identification of the catalytic sites and binding pockets which are essential for the design of novel TKIs. Therefore, multidisciplinary collaborations integrating structural biology into drug screening applications are essential to treatment of CML [20]. From this perspective, in this study, we introduce a method for the production of recombinant ABL-1 for structure-based drug screening applications. In addition, we are keen to implement these methodologies with regard to potential ABL-1 inhibitors that have been synthesized by our research group.

Our group is dedicated to the development of new bioactive substances with potential pharmaceutical applications, in particular anticancer agents carrying quinone moiety. One of the potential anticancer effects of these compounds is anti-CML activity, particularly through ABL1 inhibition [26-30]. We synthesized 2,3-dimethyl-5-amino-1,4-benzoquinones (**PQ1-15**) and determined that **PQ2** showed anti-CML activity with an IC_50_ value of 6.40 μM. PQ2 inhibited ABL1 with an IC_50_ value of 19.22 μM [26]. In other study, amino-1,4-benzoquinones (**AQ1-18**) were synthesized and investigated for anti-CML activity via ABL1 inhibition. **AQ15** induced strong cytotoxicity against K562 CML cells with an IC_50_ value of 0.76 μM and inhibited ABL1 with an IC_50_ value of 17.92 μM [27]. Besides, new chlorinated quinone analogs (**ABQ1-17**) were evaluated for their anti-CML and ABL1 inhibitory effects. **ABQ11** revealed remarkable anti-CML effects with an IC_50_ value of 0.28 μM through ABL1 inhibition (IC_50_ = 13.12 μM) [28]. Hereinafter, we synthesized quinolinequinone analogs with alkoxy group(s) in aminophenyl ring (**AQQ1-15**) and searched for their mechanistic anti-CML potential. **AQQ13** was found to be cytotoxic against CML cell line with an IC_50_ value of 0.59 μM and showed 32% ABL1 inhibition at 10 μM concentration [29]. In other study, **AQQ15** showed anti-CML activity with an IC_50_ value of 0.52 μM among new quinolinequinones (**AQQ1-19**). **AQQ15** demonstrated 35% ABL1 inhibition at 10 μM concentration [30].

This study presents an optimized method for the recombinant expression of the human ABL-1 kinase domain in *E. coli* Rosetta2™ BL21. The sarkosyl detergent was used to solubilize ABL-1 inclusion bodies that were observed under the native culture condition. After sarkosyl treatment, soluble ABL-1 was efficiently purified via Ni-NTA affinity chromatography. Following the Ni-NTA purification, 6X-His-SUMO-Tag was cleaved by ULP1 protease. The produced ABL-1 was subsequently used in crystallization studies to determine its structure via X-ray crystallography and further drug-screening applications.

## Acknowledgement

This study was supported by the Health Institutes of Türkiye (TUSEB) under the Grant Number 24333. The authors thank TUSEB for their support. Grateful acknowledgment is extended to Reyhan Kamış for her contributions to the project, especially in crystallization trials. The authors gratefully acknowledge Assoc. Prof. Belgin Sever for her insightful contributions to the tyrosine kinase inhibitor research.

